# Asymmetric transcallosal conduction delay leads to finer bimanual coordination

**DOI:** 10.1101/2020.01.24.918102

**Authors:** Marta Bortoletto, Laura Bonzano, Agnese Zazio, Clarissa Ferrari, Ludovico Pedullà, Roberto Gasparotti, Carlo Miniussi, Marco Bove

## Abstract

It has been theorized that hemispheric dominance and a more segregated information processing have evolved to overcome long conduction delay through the corpus callosum (TCD) but that this may still impact behavioral performance mostly in tasks requiring high timing accuracy. Nevertheless, a thorough understanding of the temporal features of interhemispheric communication is missing due to methodological shortcomings. Here, we show in the motor system that TCD can be measured from transcranial magnetic stimulation (TMS) -evoked potentials (TEPs): by integrating TEPs with diffusion tensor imaging (DTI) and peripheral measures of interhemispheric inhibition (i.e., the ipsilateral silent period-iSP), we show that P15 TEP component reflects TCD between motor areas. Importantly, we report that better bimanual coordination is achieved when TCD between motor areas is asymmetric. These results suggest that interhemispheric communication can be optimized through asymmetric connectivity, in which information transfer is faster from the dominant hemisphere to the non-dominant hemisphere.

## Introduction

Conduction delay over long-range connections is a crucial feature of neural communication that impacts the efficacy of signal transmission between distant areas and thus influences the anatomo-functional architecture of the brain. Specifically, long transcallosal conduction delay (TCD) has been theorized to be the basis of hemispheric dominance: long TCD prevents the exchange of information between homologous cortical areas and favors the compartmentalization of signal processing (1–3). Such delays impact each transcallosal transfer of information regardless of the information conveyed, i.e., both when the processes of the two hemispheres must be integrated and when the two hemispheres exert mutual functional inhibition, possibly directed toward suppression of competing activation, as has been shown in the motor system (4). The impact of TCD on interhemispheric signal transmission may eventually have consequences on behavioral performance, becoming most apparent when tasks have strict timing constraints (1).

Despite the acknowledged importance of TCD in brain functioning and initial indications that TCD affects cognitive functions (5, 6), empirical support has been limited to date due to the lack of a direct noninvasive measure of TCD. Pioneering studies have exploited lateralized effects on reaction times and event-related potentials, but these effects may be affected by several stages along the processing stream (7–9). In relation to the motor system, estimates of TCD have been obtained with peripheral measures of transcallosal inhibition, such as the ipsilateral silent period (iSP) (10–14), but they are affected by the corticospinal tract. Consequently, it is not well understood how conduction delay in transcallosal connections affects lateralized processing and behavioral outcomes.

Coregistration of transcranial magnetic stimulation (TMS) and electroencephalography (EEG) has the potential to provide temporally precise cortical measures of effective connectivity through TMS-evoked potentials (TEPs): After the direct activation of a target region at the time of TMS, a secondary neural response is generated in distant connected regions, e.g., a homologous area connected via the corpus callosum, and this response is recorded through EEG (15). Importantly, the amplitude and latency of the secondary response can be measured from the TEPs and reflect the strength and conduction delay of the connection, respectively.

In this work, we hypothesized that an early contralateral component of TMS-EEG coregistration could represent the response of the contralateral primary motor cortex (M1) after signal transmission through callosal fibers. In the following analyses, we show that a TEP component occurring at about 15 ms (P15) reflects transcallosal inhibitory control of the contralateral motor area; Indeed, P15 amplitude is related to inhibition of the contralateral M1 as measured by iSP. Importantly, P15 latency depends on the diffusivity of water molecules along the fibers of the callosal body, i.e., the section of the CC that connects homologous motor cortices. Therefore, P15 latency provides an index of TCD. With this new measure of effective connectivity, we tested the hypothesis that TCD impacts behavioral performance when interhemispheric activity has to be tuned with high timing accuracy. As behavioral task, we adopted a bimanual coordination task based on sequences of finger opposition movements. Previous studies have shown that time lag between hands is influenced by callosal integrity in multiple sclerosis (16) and in callosotomy and agenesis of CC (17, 18).

We show that asymmetry in TCD between motor cortices is beneficial for bimanual coordination: Specifically, shorter left-to-right TCD and longer right-to-left TCD resulted in better temporal performance in bimanual finger opposition movements. These findings suggest that, for in-phase bimanual movements, fast interhemispheric signal transmission per se (i.e., in both directions) is not as beneficial as an asymmetric interhemispheric signal transmission in which the TCD from the dominant M1 is shorter than the TCD from the nondominant M1.

## Results

In our experiment (Fig. 1), we assessed the microstructural integrity of the corpus callosum by means of diffusion tensor imaging (DTI)-derived parameters (fractional anisotropy, mean diffusivity, radial diffusivity and axial diffusivity) as well as bimanual coordination performance during in-phase bimanual sequences of thumb-to-finger opposition movements in healthy subjects (n = 15). Moreover, TEPs and iSP were collected from the left and right M1 separately during an iSP paradigm, in which the application of TMS over M1 induces a reduction in electromyographic activity of the ipsilateral target hand muscle due to transcallosal inhibition (11, 19). To increase the range of motor inhibition, we manipulated the activity of the contralateral hand by including a condition in which the hand was at rest (NoTask) and a condition in which subjects performed thumb-to-finger opposition movements (Task) (20). To account for the hierarchical structure of the design in which measures were repeated within subjects, e.g. data from Task and NoTask conditions and from left and right TMS, data were analyzed with linear mixed models.

**Fig. 1.**
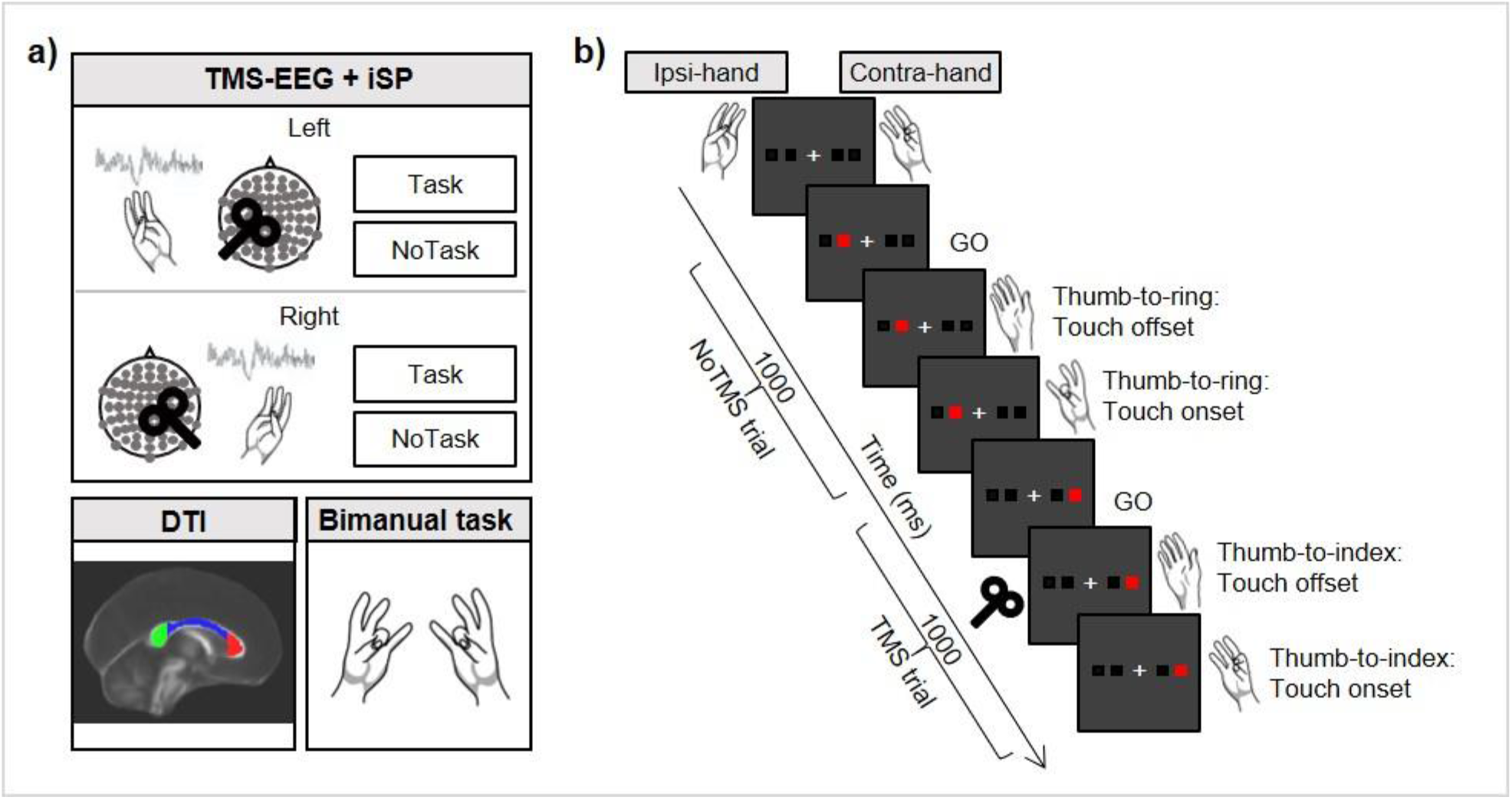
Study methods. a) Main steps of the experimental procedure, consisting of a TMS-EEG and iSP session, DTI acquisition and an in-phase bimanual coordination task. During TMS-EEG, the left and right M1 were stimulated in separate blocks in an iSP paradigm, involving Task and NoTask conditions, in counterbalanced order. During both conditions, the thumb and the little finger of the ipsilateral hand were opposed, maintaining ~25% of maximal APB muscle contraction. iSP is a reduction in electromyographic activity in the APB muscle after TMS due to transcallosal inhibition. In the NoTask condition, participants kept the contralateral hand at rest, while in the Task condition, they performed the unimanual finger opposition movement sequence described in b). b) Two example trials of the Task condition, comprising one trial without and one trial with TMS over the left M1. On the contralateral hand, the thumb was opposed to the finger indicated by the red square on the PC screen. A TMS pulse was triggered by the touch offset in half of the trials.

The stimulation of the targeted M1 induced a complex TEP response (Fig. 2a), including an early component, i.e., the abovementioned P15. The latency of ~15ms falls in the range of TCD estimated from anatomical studies (2, 21) and double-coil TMS studies (4). The peak is located in the frontocentral sites of the contralateral hemisphere. The polarity is positive, in line with the relationship between positivity and inhibition that has been shown in motor areas. Importantly, P15 was highly consistent and could be detected in every condition (Fig. 2b-c), and the same was true of the iSP (Table 1).

First, P15 was linked to contralateral motor inhibition: we found that P15 amplitude predicts the normalized iSP area (*t* = 3.33, *p* =0.001), such that the larger the P15, the stronger the inhibition will be in the ipsilateral *abductor pollicis brevis* (APB; Fig. 2d). No significant relationship was found between P15 latency and iSP onset (*t* = 1.19, *p* = 0.24).

Moreover, as evidence that P15 reflects the timing of transcallosal connectivity, we assessed whether microstructural integrity of the corpus callosum predicts the latency of P15. We expected a significant relationship for the CC body, because this section connects the two primary motor areas. We found that P15 latency was predicted by the mean diffusivity of the CC body (*t* = −2.23, *p* = 0.04) and not by its fractional anisotropy (*t* = −0.36, *p* = 0.73). Crucially, the result concerning the mean diffusivity of the callosal body was explained by the diffusivity along the axons (axial diffusivity; *t* = −2.42, *p* = 0.03) and not by the radial diffusivity (*t* = −0.89, *p* = 0.39): the higher the axial diffusivity, the shorter the latency of P15, i.e., shorter TCD (Fig. 2e). As a control, we tested that the relationship was specific for the callosal body and not for the other regions of the CC. Accordingly, no significant relationship was found for genu (*t* = −0.65, *p* = 0.53) and splenium (*t* = −0.37, *p* = 0.72).

Taken together, these results strongly support the idea that P15 reflects the transcallosal inhibition of M1 and that its latency represents the TCD along the fibers of the callosal body.

**Fig. 2.**
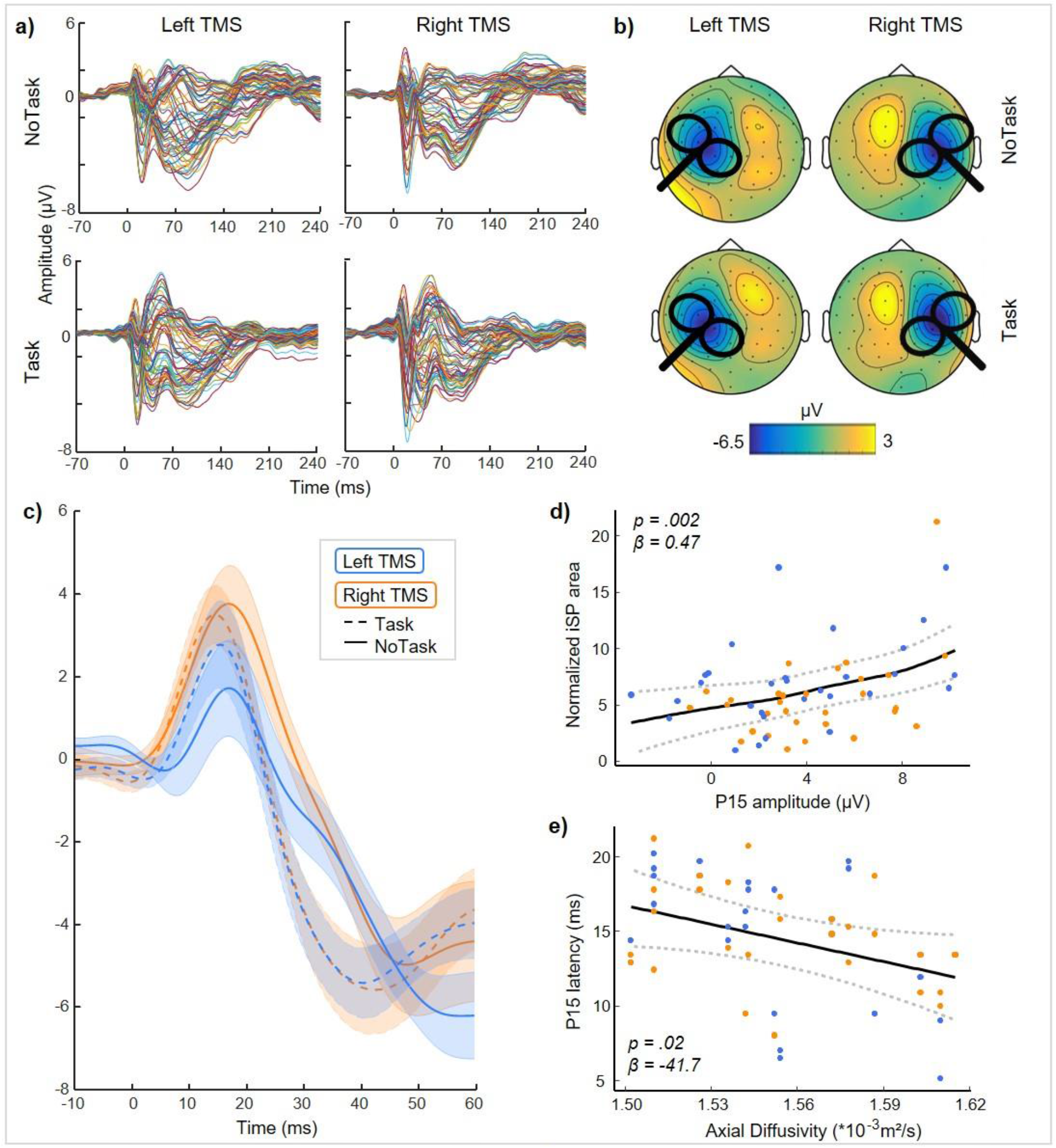
P15 as a measure of transcallosal effective connectivity. a) Grand average of TEPs in the four experimental conditions. b) Topographical maps of P15 showing a consistent pattern of positive activation in frontal electrodes contralateral to TMS in the four experimental conditions. c) Grand average of P15 in the four experimental conditions (SE in shaded error bars). P15 was identified in each participant and each condition as the first positive peak within a 5-30 ms interval in pooled data from two frontal electrodes contralateral to TMS (F1 and FC1 for right TMS, F2 and FC2 for left TMS). d) Relationship between P15 amplitude and normalized iSP area: higher P15 is associated with greater iSP, suggesting that P15 reflects transcallosal inhibition. e) Relationship between axial diffusivity in the body of the corpus callosum and P15 latency: higher axial diffusivity predicts shorter P15 latency. In d) and e), blue dots indicate left TMS, and orange dots indicate right TMS. Data from the Task and NoTask conditions were pooled together. Fitted curves were carried out through smoothed spline methods applied to predicted values-obtained by bootstrap procedure-of the LMMs.

Our next goal was to test how TCD affects behavior. Based on previous studies (16), we expected that TCD between homologous motor areas could affect the temporal precision of motor performance when bilateral movements must be coordinated. Therefore, we calculated the inter-hand interval, i.e., the time difference between taps with the right and left hand, during in-phase bimanual sequences of finger opposition movements.

We found that P15 latency for each direction of interhemispheric transfer separately predicts inter-hand interval, with opposite effects. P15 latency from left-to-right hemisphere positively predicted inter-hand interval (*t* = 2.49, *p* = 0.02; Fig. 3a), such that shorter TCD resulted in a shorter inter-hand interval, i.e., better bimanual coordination. In the opposite direction, the shorter the right-to-left P15 latency, the longer the inter-hand interval, indicating worse bimanual coordination (*t* = −2.2, *p* = 0.04; Fig. 3b). Crucially, the best predictor of bimanual coordination as indicated by Akaike Information Criterion (AIC) method (22, 23) was the ratio of the P15 latency from the dominant (left) M1 to the P15 latency from the nondominant (right) M1 (*t* = 4.17, *p* = 0.001; Fig. 3c). Finally, as a control condition, we tested the relationship between P15 latency and inter-hand interval during bimanual repetitive thumb-to-index-finger opposition movements. The corpus callosum seems to be less involved in this type of movement, as shown by a different effect induced by ipsilateral rTMS as a function of the complexity of finger motor sequences (24). In this case, the relationship between P15 latency and inter-hand interval for repetitive movements did not reach statistical significance (left TMS: *t* = 0.36, *p* = 0.72; right TMS: *t* = −1.98, *p* = 0.06; left/right TMS: *t* = 1.93, *p* = 0.07).

These data show that bimanual coordination benefits from an asymmetric TCD between homologous motor areas when signal transmission from the dominant to the nondominant hemisphere is faster than transmission in the opposite direction.

**Fig. 3.**
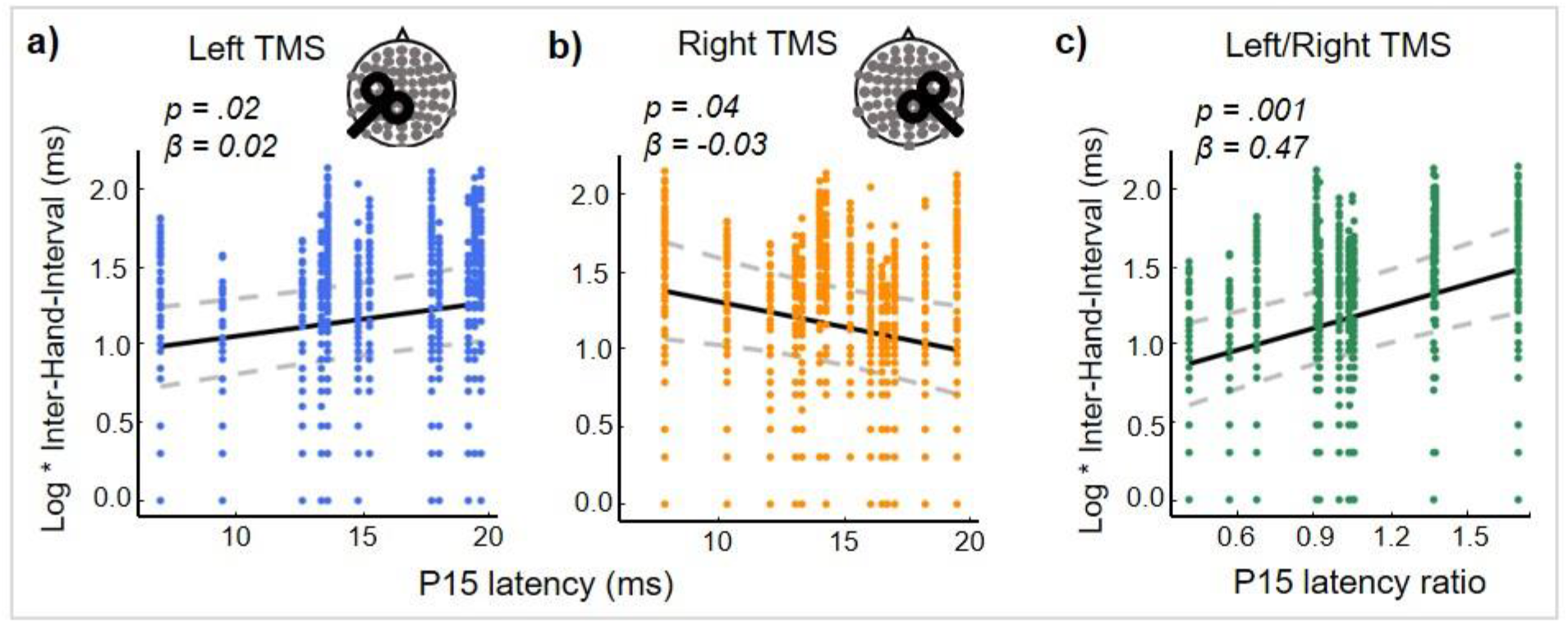
Asymmetric transcallosal conduction delay predicts finer bimanual coordination. The relationship between P15 latency and performance in the in-phase bimanual coordination task depends on the stimulated hemisphere. a) When TMS is delivered over M1 in the dominant hemisphere (left TMS), shorter P15 latency is associated with finer bimanual coordination (positive relationship between P15 latency and inter-hand interval). b) Conversely, when TMS is applied over M1 in the nondominant hemisphere (right TMS), the shorter the P15 latency is, the worse the bimanual coordination will be (negative relationship between P15 latency and inter-hand interval). c) Inter-hand interval is best predicted by the ratio of P15 latency following left TMS to P15 latency following right TMS, indicating that a shorter conduction delay from the dominant M1 to the nondominant M1 than in the opposite direction is associated with finer bimanual coordination. Fitted curves (linear trends) were carried out through smoothed spline methods applied to predicted values-obtained by bootstrap procedure-of the LMMs.

## Discussion

The present results suggest that the temporal features are crucial in the communication between hemispheres and shape the final behavioral outcome.

The temporal synchronization of bilateral movements needs an efficient interaction between the two sides of the motor system to be performed with high level of precision. According to the model of neural cross-talk, motor commands are sent from each side both to the contralateral side of the corticospinal tract and, in a mirror version, to the ipsilateral side (25–27). Pathways allowing this interaction include interhemispheric connections through the CC and subcortical pathways (28). The relative conduction delay in each direction of the CC tract may affect how the signals from the two hemispheres interact and potentially interfere with each other. In this case, a better bimanual coordination with a more efficient signal transmission from the left motor cortex is in line with the well-known dominant role of the left hemisphere in the performance of bimanual movements and in movement sequences (29–31).

Considering that P15 reflects a functionally inhibitory signal, one possible mechanism is that prompt suppression of the nondominant motor area, conveyed through the CC as a functional inhibitory signal, may increase the efficiency of cross-talk at the corticospinal level, thus improving temporal coordination. In this case, information transfer through the CC would not be necessary to perform bimanual movements but it would optimize their coordination. Accordingly, previous studies have shown that the CC contributes to temporal control of in-phase discrete movements, although CC integrity is not essential for this task, as it can be performed after callosotomy and by acallosal patients (32, 33).

Alternatively, information transfer through the CC during the bimanual task may have a facilitatory function, rather than the inhibitory function that we observed during the iSP paradigm. Therefore, the cross-talk would occur at the cortical level. This possibility cannot be ruled out because we did not record TEPs during the bimanual task. Nevertheless, faster signal transmission from the dominant hemisphere than from the nondominant hemisphere would still pose an advantage in the case of transcallosal functional facilitation, reducing the interference effects of intruding commands. Altogether, finer bimanual coordination would be reached when the transmission was asymmetric and gave a temporal advantage to the signal from the dominant hemisphere over the nondominant hemisphere, regardless of the information conveyed (i.e., either functional inhibition or signal transmission).

Furthermore, the hemispheric asymmetry in P15 latency may arise from an asymmetry in the structure of callosal connections, thus expanding the notion of transcallosal cross-talk from a functional to a structural meaning. It can be suggested that asymmetric connectivity, in which only one direction of information processing is optimized, may be a consequence of the spatial and metabolic constraints that have limited evolutionary growth of the CC relative to brain size (34–36). This optimization would improve directional information transfer from the dominant to the non-dominant hemisphere, creating the base for hemispheric dominance.

The positive relationship between P15 latency and the axial diffusivity of the callosal body is a crucial finding that supports the motion that P15 reflects the TCD. Accordingly, axial diffusivity represents the motion of water along the principal axis of the fibers rather than across it. In a healthy population, diffusivity measures may depend on several factors, including axonal diameter, myelin thickness, axon counts and density of packed fibers (37, 38). Importantly, regardless of the specific underlying anatomical characteristics, higher axial diffusivity can reflect better signal propagation.

A TEP-based estimate of TCD may be very close to the actual TCD of the fiber tract, although it may be a slight overestimate due to the time required for TMS to activate pyramidal neurons in the target region, which takes less than 1 ms (39), and the time required for activation of local circuits in the connected area, which has been estimated to be approximately 1-2 ms. Moreover, although calculating TCD based on the peak of an EEG potential has the advantage of considering the moment in which the signal-to-noise ratio is the highest, signal onset may yield a more precise calculation. Given these considerations, P15 may include an overestimation of the TCD by approximately 2-3 ms, but overall, the timing fits with the predictions of TCD derived from anatomical studies (2, 21) and from double-coil TMS studies (4).

The development of a noninvasive measure of TCD opens several new opportunities to study cortical connectivity and hemispheric asymmetries. Importantly, this approach can be extended to other cognitive domains involving other regions of the CC and other major intrahemispheric tracts. Eventually, it will be possible to integrate new knowledge on TCD in theoretical and computational models of interhemispheric interaction.

**Table 1.**
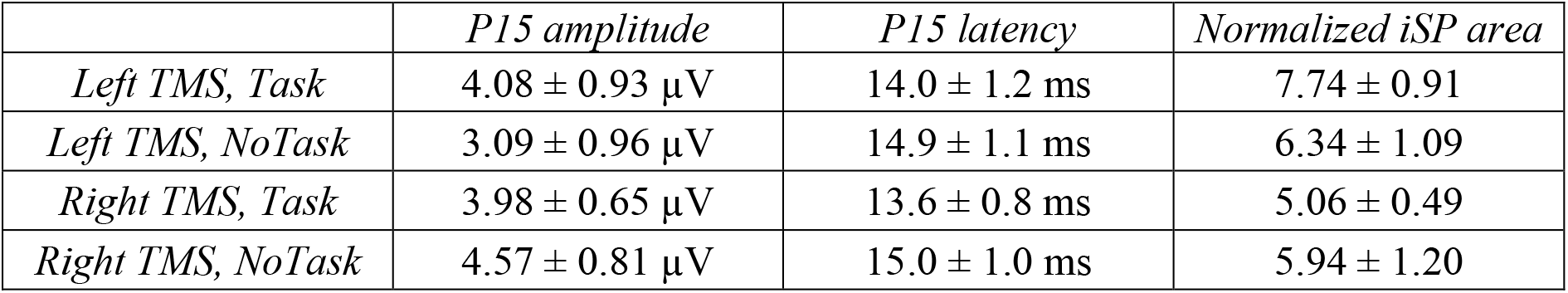
Mean ± SE Descriptive statistics (mean ± SE) of P15 amplitude, P15 latency and normalized iSP area in the four experimental conditions (Left/Right TMS X Task/NoTask)

## Methods

### Participants

Sixteen healthy participants gave written informed consent and participated in the two experimental sessions of the study within two weeks: Session 1 consisted of magnetic resonance imaging (MRI) examination, and session 2 consisted of the behavioral task and TMS-EEG for TEPs and iSP recording (Fig. 1). One participant was excluded from analyses due to technical problems during the TMS-EEG session. The final sample had mean age 35 years (range 26-47 years) and included 8 females.

All participants were right-handed according to the Edinburgh Handedness Inventory (mean ± SE: 81.5± 4.6), and they had no history of neurological disorders or contraindications to MRI or TMS. The study was performed in accordance with the ethical standards of the Declaration of Helsinki and was approved by the Ethical Committee of the IRCCS Istituto Centro San Giovanni di Dio Fatebenefratelli (Brescia) and by the Ethical Committee of the Hospital of Brescia.

### MRI acquisition

MRI was performed on a 3 T MR system (Skyra, Siemens, Erlangen, Germany). In a single session, the following scans were collected from each subject: axial T2-weighted fluid-attenuated inversion recovery (FLAIR; repetition time (TR) 9000 ms, echo time (TE) 76 ms, inversion time (TI) 2500 ms, slice thickness 3 mm, distance factor 10%, 1 average, field of view (FOV) 220 mm, voxel size 0.6×0.6×3.00 mm), DTI with spin-echo echo-planar axial sequences (multiband, TR 4100 ms, TE 75.0 ms, 1.8 mm isotropic resolution, b 1000 s/mm^2^, 64 encoding directions, 5 b0 images, fat suppression), and high-resolution T1-weighted 3D anatomical sequences (sagittal volume, TR 2400 ms, TE 2 ms, 0.9 mm isotropic resolution).

### Bimanual coordination task

Participants were seated in a comfortable chair in a quiet room, resting their forearms on a table, and were asked to perform an in-phase bimanual task before the TMS-EEG session. The task consisted of performing repetitive, metronome-paced thumb-to-finger opposition movements at 2 Hz with their eyes closed; participants performed the task with both hands simultaneously to assess bimanual coordination (16). The motor sequences consisted of simple finger tapping (thumb-to-index-finger opposition) and a 4-item sequence that consisted of opposing the thumb to the index, middle, ring and little fingers. Each condition was performed twice in separate trials lasting 45 s and separated by a few minutes of rest to avoid fatigue effects. Finger contacts were recorded by two specially designed gloves (GAS, ETT, s.p.a., Genoa, Italy) (40–42).

### TMS-EEG acquisition

Participants were comfortably seated in a dimly lit room in front of a computer screen, wearing an EEG cap and two gloves with integrated sensors. The participants were asked to perform two conditions (Task and NoTask) of an iSP paradigm in separate blocks while TMS-EEG was recorded. In the hand ipsilateral to the stimulation, the thumb and the little finger were opposed and contracted in both conditions (mean ± SE of percentage of maximal contraction: Task condition, 23% ± 1; NoTask condition, 23% ± 2). The activity in the hand contralateral to the stimulation depended on the condition. In the Task condition (Fig. 1b), the contralateral hand performed a unimanual finger tapping task. Participants were presented with four white squares on the distal phalanges of the index, middle, ring and little fingers. The white squares turned red one at a time in random order, and participants were instructed to respond as quickly and accurately as possible by opposing the thumb to the corresponding finger. The block started with participants in a resting position, touching the tip of the index finger to the tip of the thumb. Upon the presentation of the stimuli, participants lifted their fingers (touch offset) and tapped their thumb to the finger indicated by the stimulus (touch onset). Stimuli lasted 1000 ms and were presented at a frequency of 1 Hz. The number of stimuli per block was 120. Before the beginning of the recording, participants performed one block of the task with each hand to familiarize them with the task and to measure their reaction times (touch offset). Performance was not further analyzed in those blocks.

In the NoTask condition, participants saw the same stimuli as in the Task condition, but they were not required to perform any tapping with the contralateral hand, which was relaxed.

TMS over the M1 was randomly delivered in half of the trials, i.e., 60 pulses per block, at the time of touch offset measured by the engineered glove in the Task condition, and at the time of touch offset measured in the training block for the NoTask condition.

The stimulation was performed with a MagPro X100 including MagOption (MagVenture, Denmark) and set to deliver biphasic single pulses with a figure-of-eight C-B60 coil. The recharge delay was set at 500 ms. The coil was positioned tangentially to the scalp over the M1 hotspot, which was functionally localized as the position that induced reliable motor evoked potentials (MEPs) in the APB. The coil, with the handle pointing backward, was rotated away from the midline by approximately 45° so that the current induced in the cortex followed the optimal direction, i.e., anterior to posterior and posterior to interior (AP-PA). The stimulation intensity (mean ± SE: 58.1% of MSO ± 1.6%) was set at 110% of the individual average resting motor threshold (rMT), defined as the minimum TMS intensity to elicit an MEP of at least 50 μV in 5 out of 10 trials (43).

In order to ensure the precision of stimulation, a stereotaxic neuronavigation system (SofTaxic, EMS, Italy) was used in which the T1 anatomical MRI was coregistered to head position.

EEG was recorded with a TMS-compatible EEG system (BrainAmp, Brain Products GmbH, Munich, Germany) from 67 channels according to the international 10-20 system (sampling rate: 5 kHz; online bandpass filter: between 0.1 and 1 kHz). The ground was placed at FPz, and all channels were referenced online to the nose. The skin/electrode impedance was below 5 kΩ. Vertical and horizontal eye movements were monitored with an electrooculogram using two pairs of electrodes in a bipolar montage. Electromyography (EMG) was recorded from the APBs of both hands using a pair of surface electrodes with a belly-tendon montage. Before TMS-EEG, EMG was recorded for 30 s while participants were asked to touch the little finger to the thumb and to maintain the muscle contraction at maximum strength. This recording was subsequently analyzed to calculate the relative contraction levels during TMS-EEG.

### DTI analysis

DTI data were processed using FMRIB’s Diffusion Toolbox (FDT) (44). After correction for eddy current distortions and motion artifacts, a diffusion tensor model was fitted at each voxel, and the three eigenvalues were calculated (45). Parametric maps were obtained for fractional anisotropy, mean diffusivity, axial diffusivity (i.e., water diffusivity parallel to the axonal fibers), and radial diffusivity (i.e., water diffusivity perpendicular to the axonal fibers) (38, 46). All these maps were then nonlinearly transformed and aligned to 1 × 1 × 1 mm standard space using tract-based spatial statistics (TBSS) routines (47). The mean value of each DTI-derived parameter was calculated for each scan in the voxels included in the callosal fibers within three ROIs (genu, body, and splenium) from the JHU ICBM 81 white matter label atlas included in FSL (48).

### Bimanual coordination assessment

To quantitatively evaluate bimanual coordination performance, we calculated the inter-hand interval for each tap as time difference between the onset of a finger tap with the left hand and the onset of the corresponding finger tap with the right hand and removed inter-hand interval values that were greater than two standard deviations from the mean. Then, we calculated the absolute value for each tap: the longer the inter-hand interval value, the worse the bimanual coordination (16). Finally, we log-transformed the data to obtain normal distribution. Data from one participant were missing due to technical problems to the gloves.

### TMS-evoked potentials (TEPs)

TMS-EEG data analysis was performed in MATLAB (The MathWorks, Natick, MA, USA) with custom scripts using EEGLAB functions (49), FieldTrip functions (50), the source-estimate-utilizing noise-discarding (SOUND) algorithm (51) and the signal-space projection and source-informed reconstruction (SSP-SIR) algorithm (52). Continuous EEG was linearly interpolated from 1 ms before to 6 ms after the TMS pulse and high-pass filtered at 0.1 Hz. TMS-EEG data were then epoched from −200 ms before to 500 ms after TMS and downsampled to 2048 Hz. Measurement noise was discarded with the SOUND algorithm with the same spherical 3-layer model and regularization parameter (λ = 0.01) described in the original work (51). After the application of the SOUND algorithm, the signal was visually inspected, and initial artifact rejection was performed; then, independent component analysis (ICA; infomax algorithm) was run to correct ocular artifacts. TMS-evoked muscular artifacts in the first 50 ms were removed using SSP-SIR, a method based on signal-space projection and source-informed reconstruction. Muscle-artifact components (0-3 in each dataset) were identified from the time-frequency pattern and corresponding signal power. Then, epochs were low-pass filtered at 70 Hz and re-referenced to the average of TP9 and TP10. Finally, after a second visual inspection and artifact rejection, TMS-EEG data were baseline corrected from −100 ms to −2 ms before the TMS pulse and averaged. P15 amplitude and latency were measured by identifying each individual subject’s first positive peak between 5 and 30 ms in pooled data from two frontocentral channels (F2-FC2 for left TMS, F1-FC1 for right TMS).

### Ipsilateral silent period (iSP)

iSP parameters were assessed in the trace obtained from averaging the 60 rectified EMG traces (11). The following iSP parameters were considered: the iSP onset, defined as the point after cortical stimulation at which EMG activity became constantly (for a minimum duration 10 ms) below the mean amplitude of EMG activity preceding the cortical stimulus; the iSP duration, calculated by subtracting the onset time from the ending time (i.e., the first point after iSP onset at which the level of EMG activity returned to the mean EMG signal); and the normalized iSP area, calculated using the following formula: [(area of the rectangle defined as the mean EMG × iSP duration)-(area underneath the iSP)] divided by the EMG signal preceding the cortical stimulus.

### Statistical analysis

Relationships between variables were tested by linear mixed models (LMMs) with random slope and intercept (53). A summary is reported in table S1 in Supplementary materials. Akaike information Criterion (AIC) was used to find the best predictors in terms of model goodness of fit.

To test the relationship between P15 amplitude (repeated independent variable) and iSP normalized area (repeated dependent variable), a LMM was run with Condition (4 levels: Task, NoTask, LeftTMS, RightTMS) and Subjects as fixed and random effects, respectively, with each condition repeated within Subjects. The same model was employed to evaluate the relationship between P15 latency and iSP onset.

To study the predictive value of microstructural integrity of CC body (i.e., DTI measures as independent variables) on P15 latency (dependent variable), two separate LMMs were applied for fractional anisotropy and mean diffusivity of CC body. Moreover, we tested the direction of diffusivity in the CC body that predicted P15 latency by employing axial or radial diffusivity as predictors in two separate LMMs. In addition, mean diffusivity for other regions of the CC was evaluated by carrying out other two LMMs with P15 latency as dependent variable and CC genu and CC splenium as predictors.

Finally, we tested the relationship between P15 latency (as predictor) and bimanual coordination performance (i.e., inter-hand interval, as dependent variable) in a sequential thumb-to-finger opposition movement task. Separate LMMs were performed considering the three measures of P15 latency (i.e. mean value of Task and NoTask condition in: Left TMS, Right TMS and the rate between the two) as predictors, each tap (Tap) of the bimanual task as fixed effect and Subjects as random effect (with Tap repeated within Subjects). The same models were run for the repetitive thumb-to-index finger opposition movements task.

The predicted values (and corresponding standard error) of the LMMs were obtained by bootstrap procedure (number of simulation n=500). Fitted curves of predicted values were carried out through smoothed spline methods.

Statistical significance was set at p < 0.05. All the analyses were performed in R software (R Core Team (2013), http://www.R-project.org/.) and LMM were estimated by lme4 R package.

## Acknowledgements

This work was supported by the Italian multiple sclerosis foundation (FISM) Grant 2016/R/2. We would like to thank Alice Bollini and Simona Finazzi for their support with data acquisition.

## Contributions

M. Bortoletto, L.B., R.G., C.M. and M. Bove designed the research. M. Bortoletto, L.B. and R.G. performed experiments. M. Bortoletto, L.B., A.Z., L.P., C.F. performed data analyses. M. Bortoletto, L.B., A.Z., C.M., M. Bove wrote the manuscript. All authors critically reviewed data and edited the final manuscript.

## Ethics declarations

Competing interests

The authors declare no competing interests.

## Data availability

The data that support the findings of this study are available from the authors on reasonable request.

---

**Table S1.** Structure and results of LMMs. All models include subjects as random factor. *β*, *t* and *p* values refer to the independent variable. Asterisks indicates significant effects (*p* < 0.05).

| | Dependent variable | Independent variable | Fixed factor repeated within Subjects | $\beta$ | $t$ | $p$ | AIC |
| --- | --- | --- | --- | --- | --- | --- | --- |
| *I) P15 amplitude and iSP* |  |  |  |  |  |  |  |
| Ia) | Normalized iSP area | P15 amplitude | Condition (4 measures per subject) | 0.47 | 3.33 | 0.002 * | 326.7 |
| Ib) | iSP onset | P15 latency | Condition (4 measures per subject) | 195.11 | 1.19 | 0.24 | 375.7 |
| *II) CC microstructural integrity and P15 latency* |  |  |  |  |  |  |  |
| IIa) | P15 latency | Fractional anisotropy of CC body | Condition (4 measures per subject) | -0.02 | -0.36 | 0.73 | -500.7 |
| IIb) | P15 latency | Mean diffusivity of CC body | Condition (4 measures per subject) | -68.58 | -2.23 | 0.04 * | -504.2 |
| IIc) | P15 latency | Mean diffusivity of CC splenium | Condition (4 measures per subject) | -6.13 | -0.37 | 0.72 | -500.7 |
| IId) | P15 latency | Mean diffusivity of CC genu | Condition (4 measures per subject) | -22.2 | -0.65 | 0.53 | -501.0 |
| IIe) | P15 latency | Axial diffusivity of CC body | Condition (4 measures per subject) | -41.66 | -2.42 | 0.02 * | -505.4 |
| IIf) | P15 latency | Radial diffusivity of CC body | Condition (4 measures per subject) | -28.48 | -0.89 | 0.39 | -501.2 |
| *III) P15 latency and bimanual coordination* |  |  |  |  |  |  |  |
| IIIa) | Inter-hand interval (thumb-to-finger sequence) | P15 amplitude (left TMS) | Tap (91 measures per subject) | 0.02 | 2.49 | 0.02 * | 1605.7 |
| IIIb) | Inter-hand interval (thumb-to-finger sequence) | P15 amplitude (right TMS) | Tap (91 measures per subject) | -0.03 | -2.2 | 0.04 * | 1603.3 |
| IIIc) | Inter-hand interval (thumb-to-finger sequence) | P15 amplitude (left/right TMS) | Tap (91 measures per subject) | 0.47 | 4.17 | 0.001 * | 1596.4 |
| IIId) | Inter-hand interval (thumb-to-index repetitions) | P15 amplitude (left TMS) | Tap (81 measures per subject) | 0.003 | 0.36 | 0.72 | 1264.6 |
| IIIe) | Inter-hand interval (thumb-to-index repetitions) | P15 amplitude (right TMS) | Tap (81 measures per subject) | -0.02 | -1.98 | 0.06 | 1261.8 |
| IIIf) | Inter-hand interval (thumb-to-index repetitions) | P15 amplitude (left/right TMS) | Tap (81 measures per subject) | 0.19 | 1.93 | 0.07 | 1260.7 |

## References

1. J. L. Ringo, R. W. Doty, S. Demeter, P. Y. Simard, Time Is of the Essence: A Conjecture That Hemispheric Specialization Arises From Interhemispheric Conduction Delay. Cereb. Cortex 4, 331–43 (1994).

2. K. A. Phillips, et al., The corpus callosum in primates : processing speed of axons and the evolution of hemispheric asymmetry. Proc. R. Soc. B Biol. Sci. 282, 20151535 (2015).

3. V. R. Karolis, M. Corbetta, M. T. De Schotten, The architecture of functional lateralisation and its relationship to callosal connectivity in the human brain. Nat. Commun. 10, 1417 (2019).

4. A. Ferbert, et al., Interhemispheric Inhibition of the Human Motor Cortex. J. Physiol. 453, 525–546 (1992).

5. M. J. Hoptman, R. J. Davidson, How and why do the two cerebral hemispheres interact? Psychol. Bull. 116, 195–219 (1994).

6. N. Cherbuin, C. Brinkman, Efficiency of callosal transfer and hemispheric interaction. Neuropsychology 20, 178–184 (2006).

7. C. A. Marzi, P. Bisiacchi, R. Nicoletti, Is interhemispheric transfer of visuomotor information asymmetric? Evidence from a meta-analysis. Neuropsychologia 29, 1163–1177 (1991).

8. R. Chaumillon, J. Blouin, A. Guillaume, Eye dominance influences triggering action: The Poffenberger paradigm revisited. Cortex 58, 86–98 (2014).

9. R. Chaumillon, J. Blouin, A. Guillaume, Interhemispheric transfer time asymmetry of visual information depends on eye dominance: An electrophysiological study. Front. Neurosci. 12, 1–19 (2018).

10. T. Davidson, F. Tremblay, Hemispheric Differences in Corticospinal Excitability and in Transcallosal Inhibition in Relation to Degree of Handedness. PLoS One 8, 1–9 (2013).

11. C. Trompetto, M. Bove, L. Marinelli, L. Avanzino, A. Buccolieri, Suppression of the transcallosal motor output: a transcranial magnetic stimulation study in healthy subjects. Exp. brain Res. 158, 133–140 (2004).

12. B. W. Fling, B. L. Benson, R. D. Seidler, Transcallosal sensorimotor fiber tract structure-function relationships. Hum. Brain Mapp. 34, 384–395 (2013).

13. B. U. Meyer, S. Röricht, C. Woiciechowsky, Topography of fibers in the human corpus callosum mediating interhemispheric inhibition between the motor cortices. Ann. Neurol. 43, 360–369 (1998).

14. B. U. Meyer, S. Röricht, H. G. Von Einsiedel, F. Kruggel, A. Weindl, Inhibitory and excitatory interhemispheric transfers between motor cortical areas in normal humans and patients with abnormalities of the corpus callosum. Brain 118, 429–440 (1995).

15. M. Bortoletto, D. Veniero, G. Thut, C. Miniussi, The contribution of TMS – EEG coregistration in the exploration of the human cortical connectome. Neurosci. Biobehav. Rev. 49, 114–124 (2015).

16. L. Bonzano, et al., Callosal Contributions to Simultaneous Bimanual Finger Movements. J. Neurosci. 28, 3227–3233 (2008).

17. C. Ouimet, et al., Bimanual crossed-uncrossed difference and asynchrony of normal, anterior- and totally-split-brain individuals. Neuropsychologia 48, 3802–3814 (2010).

18. J. C. Eliassen, K. Baynes, M. S. Gazzaniga, Anterior and posterior callosal contributions to simultaneous bimanual movements of the hands and fingers. Brain 123, 2501–2511 (2000).

19. M. Wahl, et al., Human motor corpus callosum: Topography, somatotopy, and link between microstructure and function. J. Neurosci. 27, 12132–12138 (2007).

20. F. Giovannelli, et al., Modulation of interhemispheric inhibition by volitional motor activity: An ipsilateral silent period study. J. Physiol. 587, 5393–5410 (2009).

21. R. Caminiti, et al., Diameter, length, speed, and conduction delay of callosal axons in macaque monkeys and humans: Comparing data from histology and magnetic resonance imaging diffusion tractography. J. Neurosci. 33, 14501–14511 (2013).

22. K. P. Burnham, D. R. Anderson, Model Selection and Multimodel Inference: A Practical Information-Theoretic Approach (2nd ed) (2002).

23. K. P. Burnham, D. R. Anderson, Multimodel inference: Understanding AIC and BIC in model selection. Sociol. Methods Res. 33, 261–304 (2004).

24. L. Avanzino, et al., 1-Hz repetitive TMS over ipsilateral motor cortex influences the performance of sequential finger movements of different complexity. 27, 1285–1291 (2008).

25. D. Cattaert, A. Semjen, J. J. Summers, Simulating a neural cross-talk model for between-hand interference during bimanual circle drawing. Biol. Cybern. 358, 343–358 (1999).

26. Y. Aramaki, M. Honda, N. Sadato, Suppression of the non-dominant motor cortex during bimanual symmetric finger movement: a functional magnetic resonance imaging study. Neuroscience 141, 2147–2153 (2006).

27. U. Ziemann, M. Hallett, Hemispheric asymmetry of ipsilateral motor cortex activation during unimanual motor tasks : further evidence for motor dominance. Clin. Neurophysiol. 112, 107–113 (2001).

28. R. B. Ivry, E. Hazeltine, Subcortical locus of temporal coupling in the bimanual movements of a callosotomy patient (1999).

29. G. M. Geffen, D. L. Jones, L. B. Geffen, Interhemispheric control of manual motor activity. Behav. Brain Res. 64, 131–140 (1994).

30. D. J. Serrien, R. B. Ivry, S. P. Swinnen, Dynamics of hemispheric specialization and integration in the context of motor control. 7, 160–167 (2006).

31. L. M. Rueda-Delgado, et al., Understanding bimanual coordination across small time scales from an electrophysiological perspective. Neurosci. Biobehav. Rev. 47, 614–635 (2014).

32. S. W. Kennerley, J. Diedrichsen, E. Hazeltine, A. Semjen, R. B. Ivry, Callosotomy patients exhibit temporal uncoupling during continuous bimanual movements. Nat. Neurosci. 5, 376–381 (2002).

33. J. Gooijers, S. P. Swinnen, Interactions between brain structure and behavior: The corpus callosum and bimanual coordination. Neurosci. Biobehav. Rev. 43, 1–19 (2014).

34. R. Caminiti, H. Ghaziri, R. Galuske, P. R. Hof, G. M. Innocenti, Evolution amplified processing with temporally dispersed slow neuronal connectivity in primates. PNAS 106, 19551–6 (2009).

35. S. Herculano-Houzel, B. Mota, P. Wong, J. H. Kaas, Connectivity-driven white matter scaling and folding in primate cerebral cortex. Proc. Natl. Acad. Sci. U. S. A. 107, 19008–19013 (2010).

36. W. D. Hopkins, M. Misiura, S. M. Pope, E. M. Latash, Behavioral and brain asymmetries in primates: A preliminary evaluation of two evolutionary hypotheses. Ann. N. Y. Acad. Sci. 1359, 65–83 (2015).

37. F. Aboitiz, A. B. Scheibel, R. S. Fisher, E. Zaidel, Fiber composition of the human corpus callosum. Brain Res. 598, 143–153 (1992).

38. C. Beaulieu, The Biological Basis of Diffusion Anisotropy (Elsevier Inc., 2009) https:/doi.org/10.1016/B978-0-12-374709-9.00006-7.

39. J. K. Mueller, et al., Simultaneous transcranial magnetic stimulation and single-neuron recording in alert non-human primates. Nat. Neurosci. 17, 1130–1136 (2014).

40. A. Signori, et al., Quantitative assessment of finger motor performance : Normative data. PLoS One 12, e0186524 (2017).

41. M. Bove, et al., The effects of rate and sequence complexity on repetitive finger movements. Brain Res. 1153, 84–91 (2007).

42. L. Bonzano, et al., Quantitative Assessment of Finger Motor Impairment in Multiple Sclerosis. PLoS One 8, 1–7 (2013).

43. S. Rossi, M. Hallett, P. M. Rossini, A. Pascual-Leone, Safety, ethical considerations, and application guidelines for the use of transcranial magnetic stimulation in clinical practice and research. Clin. Neurophysiol. 120, 2008–2039 (2009).

44. S. M. Smith, et al., Advances in functional and structural MR image analysis and implementation as FSL. Neuroimage 23, 208–219 (2004).

45. B. P.J., Inferring Microstructural Features and the Physiological State of Tissues from Diffusion Weighted Images. NMR Biomed. 8, 333–344 (1995).

46. L. Bonzano, et al., NeuroImage Upper limb motor rehabilitation impacts white matter microstructure in multiple sclerosis. Neuroimage 90, 107–116 (2014).

47. S. M. Smith, et al., Tract-based spatial statistics: Voxelwise analysis of multi-subject diffusion data. Neuroimage 31, 1487–1505 (2006).

48. S. Mori, S. Wakana, P. C. M. Van Zijl, L. M. Nagae-Poetscher, MRI atlas of human white matter (Elsevier, 2005).

49. A. Delorme, S. Makeig, EEGLAB: an open source toolbox for analysis of single-trial EEG dynamics including independent component analysis. J. Neurosci. Methods 134, 9–21 (2004).

50. R. Oostenveld, P. Fries, E. Maris, J. M. Schoffelen, FieldTrip: Open source software for advanced analysis of MEG, EEG, and invasive electrophysiological data. Comput. Intell. Neurosci. 2011 (2011).

51. T. P. Mutanen, J. Metsomaa, S. Liljander, R. J. Ilmoniemi, NeuroImage Automatic and robust noise suppression in EEG and MEG : The SOUND algorithm. Neuroimage 166, 135–151 (2018).

52. T. P. Mutanen, et al., NeuroImage Recovering TMS-evoked EEG responses masked by muscle artifacts. Neuroimage 139, 157–166 (2016).

53. A. Galecki, T. Burzykowski, Linear Mixed-Effects Models Using R: A Step-by-Step Approach (Springer-Verlag New York, 2013) https:/doi.org/10.1007/978-1-4614-3900-4.

